# A haploid wild yeast resource for exploring the natural ecology of *Saccharomyces cerevisiae*

**DOI:** 10.64898/2026.02.13.705822

**Authors:** Cheng-Ju Yang, Yu-Chen Yeh, Chen Hsiao, Yu-Hsiu Liu, Min R Lu, Yu-Ching Liu, Gianni Liti, Isheng Jason Tsai, Huei-Mien Ke

## Abstract

*Saccharomyces cerevisiae* predominantly exists as diploid cells in nature, a life-cycle feature that limits classical genetic analyses of wild populations. Here, we establish a stable haploid strain collection derived from 33 diverse Taiwanese *S. cerevisiae* isolates through targeted disruption of the HO endonuclease gene. This resource spans predomesticated Asian wild lineages and enables systematic analyses of mating compatibility, reproductive isolation, and ecological trait variation. Although all pairwise hybridizations formed zygotes, many produced low spore viabilities, revealing strong postzygotic barriers. Genome analyses show that reduced hybrid fertility is primarily associated with chromosomal inversions and inter-chromosomal rearrangements rather than sequence divergence, indicating that structural variation maintains lineage separation despite geographic coexistence. Phenotypic profiling uncovered marked ecological differentiation, with the most diverged TW1 lineage favoring cooler growth conditions and a naturally occurring hybrid exhibiting heterosis with expanded thermal tolerance. While most wild strains grew poorly on maltose, two anthropogenically associated strains displayed enhanced maltose utilization linked to functional *MAL*⁺ regulatory alleles and maltose-specific transporters. Together, these findings demonstrate how structural genomic variation and metabolic gene divergence drive ecological and reproductive divergence in wild *S. cerevisiae* and establish this haploid collection as a platform for studying yeast evolution in nature.

## Introduction

*Saccharomyces cerevisiae* is known as an invaluable model organism for genetics, biotechnology, and evolutionary studies due to its relatively small genome size, extensive genetic toolkit, rapid growth rate, and historical significance in human-driven fermentations^1–7^. Laboratory-based strains, particularly S288C and derivatives, have dominated genetic and functional genomic research for decades, providing critical insights into fundamental biological processes such as cell cycle regulation, DNA repair, and metabolism. However, these domesticated strains derived from a limited number of phylogenetic lineages adapted to human-associated environment, and represent only a fraction of the vast genetic and phenotypic diversity present in wild yeast populations^8–10^. Wild isolates help uncover natural variation that may be masked in domesticated strains shaped by human-driven selection and tolerated loss-of-function mutations^11–13^. Recent advances in genomic technologies have begun to illuminate the tremendous genetic variability of wild populations, demonstrating significant divergence from classical laboratory models^14–16^. Moreover, the use of wild yeast in brewing facilitates the diversification of beer flavors, thereby enhancing beer characteristics and expanding potential brewing capabilities^17,18^.

Natural populations of *S. cerevisiae* are typically found in diploid states^14,16^, driven primarily by their homothallic nature. The homothallic lifestyle, governed by the *HO* endonuclease gene responsible for mating-type switching, ensures rapid autodiploidization of haploid spores^19,20^, severely limiting the stable existence of haploid lineages in natural environments. This inherent genetic feature poses considerable challenges for classical genetic studies, which require stable haploid strains to facilitate genetic crosses, mapping studies, and precise allele manipulations^21^. To circumvent these challenges, systematic inactivation of the *HO* gene has emerged as a powerful strategy, allowing stable propagation and controlled breeding of wild-derived haploid strains. Targeted *HO* disruption, accomplished by various molecular techniques including CRISPR-Cas9-mediated genome editing, enables the maintenance of stable haploid populations by preventing mating-type switching and subsequent autodiploidization^22,23^. This methodological breakthrough has facilitated the generation of extensive haploid strain collections, dramatically expanding the scope of genetic and evolutionary analyses beyond traditional laboratory yeast strains.

Over the past decade, several landmark studies have demonstrated the value of haploid resource collections derived from diverse wild isolates. Cubillos et al.^24^ released a seminal set of genetically tractable haploid and diploid strains from wild isolates, paving the way for comprehensive genomic and phenotypic investigations of natural yeast diversity. Such resources have enabled detailed investigations into mating compatibility and reproductive isolation mechanisms within species, uncovering various forms of post-zygotic isolation including chromosomal rearrangements and sequence divergence-driven meiotic incompatibilities^25,26^. In addition, spore viability between strains can be important for applications such as strain improvement and genetic engineering, where the success of crossbreeding or genetic manipulation may depend on the ability of different strains to mate and produce viable progeny.

Taiwanese forests represent an exceptional natural laboratory for exploring these phenomena due to their rich and diverse *S. cerevisiae* populations^14^. Multiple genetically distinct lineages coexist geographically, yet surprisingly, hybrids between these lineages remain rare. This pattern suggests the presence of strong intrinsic reproductive barriers, despite opportunities for gene flow. Establishing stable haploid lineages from these populations enables direct genetic crosses, facilitating the identification and characterization of specific genetic incompatibilities that underpin reproductive isolation. Beyond reproductive biology, stable haploid resources derived from wild isolates have proven invaluable for examining the ecological adaptations and phenotypic diversity of yeast. Wild populations exhibit remarkable variation in critical ecological traits, including thermotolerance, stress responses, and metabolic versatility across various carbon sources. For instance, Abrams et al.^27^ highlighted extensive intra-specific variation in thermotolerance, tracing significant trait differences back to specific regulatory and structural genomic variations. Similarly, comprehensive analyses of wild isolates have identified pronounced variability in the ability to metabolize alternative carbon sources, reflecting specialized adaptations to their distinct ecological niches^21^. These detailed phenotypic assessments, coupled with genomic analyses facilitated by haploid strain collections, have significantly expanded our understanding of the genetic architecture underlying ecological specialization in yeast. Such insights are crucial for comprehending how natural yeast populations adapt to varying environmental pressures, illuminating the evolutionary processes shaping phenotypic diversity.

In this study, we developed a stable haploid strain collection comprising 33 Taiwanese *S. cerevisiae* isolates representing ten genetically distinct wild lineages, including three strains associated with Asian fermentation practices. Using CRISPR-Cas9-mediated disruption^28,29^ of the *HO* endonuclease gene, we generated tractable haploids and performed systematic intraspecific crosses to assess mating compatibility, reproductive isolation, and sporulation efficiency across lineages. We also performed comprehensive phenotypic profiling, focusing on thermotolerance and carbon source utilization, to uncover ecological differentiation among lineages and hybrid strains. By integrating classical yeast genetics with the natural genomic and ecological diversity of wild populations, this resource offers a powerful platform for dissecting the genetic basis of adaptation, reproductive barriers, and phenotypic diversity in *S. cerevisiae*—with broad relevance to evolutionary biology and biotechnology.

## Results

### Generation of haploid wild Taiwanese *S. cerevisiae* strains

To establish a comprehensive haploid resource of wild *S. cerevisiae* from our extensive previous sampling efforts across Taiwan^14^, we selected 33 representative strains out of possible 121 isolates (**Fig. 1a and Table 1**). These selected strains were isolated from geographically distinct locations and diverse ecological niches, including tree-associated substrates (leaves, twigs, bark, fruit, fruit abscission, litter) and traditional fermentation products (**Table 1**). Each selected isolate represents a distinct lineage previous identified through genomic analyses^14^. Across these strains, the median pairwise genetic divergence was 17.6 SNPs per kilobase, and genome-wide heterozygosity between homologous chromosomes ranged from 1.59 × 10^−4^ to 8.94 × 10⁻^4^. Despite the coexistence of distinct lineages, no clear relationship between genetic divergence and geographic proximity was observed (**Fig. 1b**). Thus, our wild isolates, including natural hybrids, provide a valuable resource to investigate the ecological interactions and evolutionary processes driving genetic diversification in *S. cerevisiae*.

**Figure 1.**
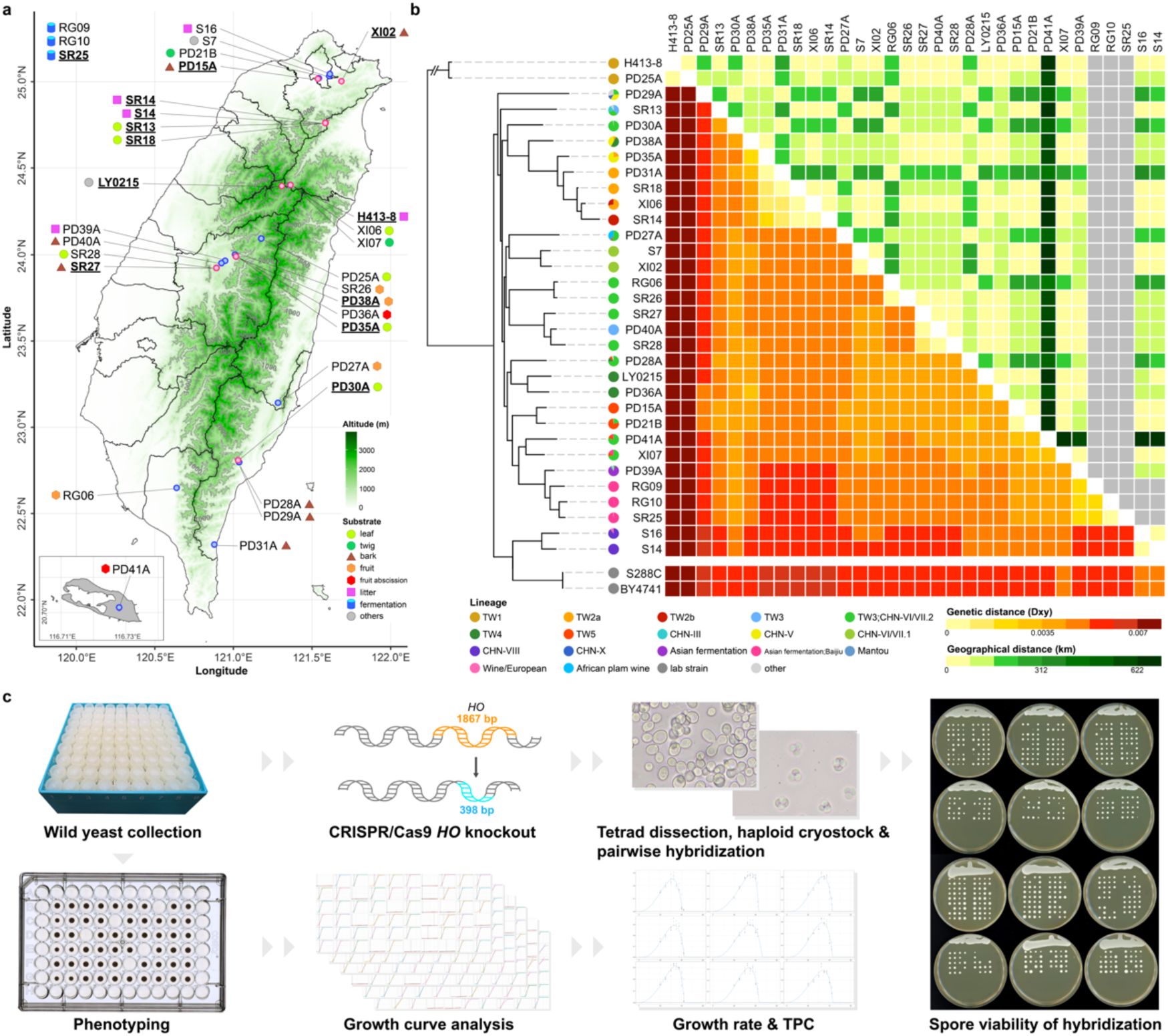
Sample description and experiment overview. **a.** Sampling locations and isolation substrates. Strains shown in bold and underlined were selected for further analysis. **b.** Comparison between pairwise genetic (bottom left) and geographical (top right) distances among isolated strains. **c.** Schematic diagram of experiment workflow.

**Table 1.**
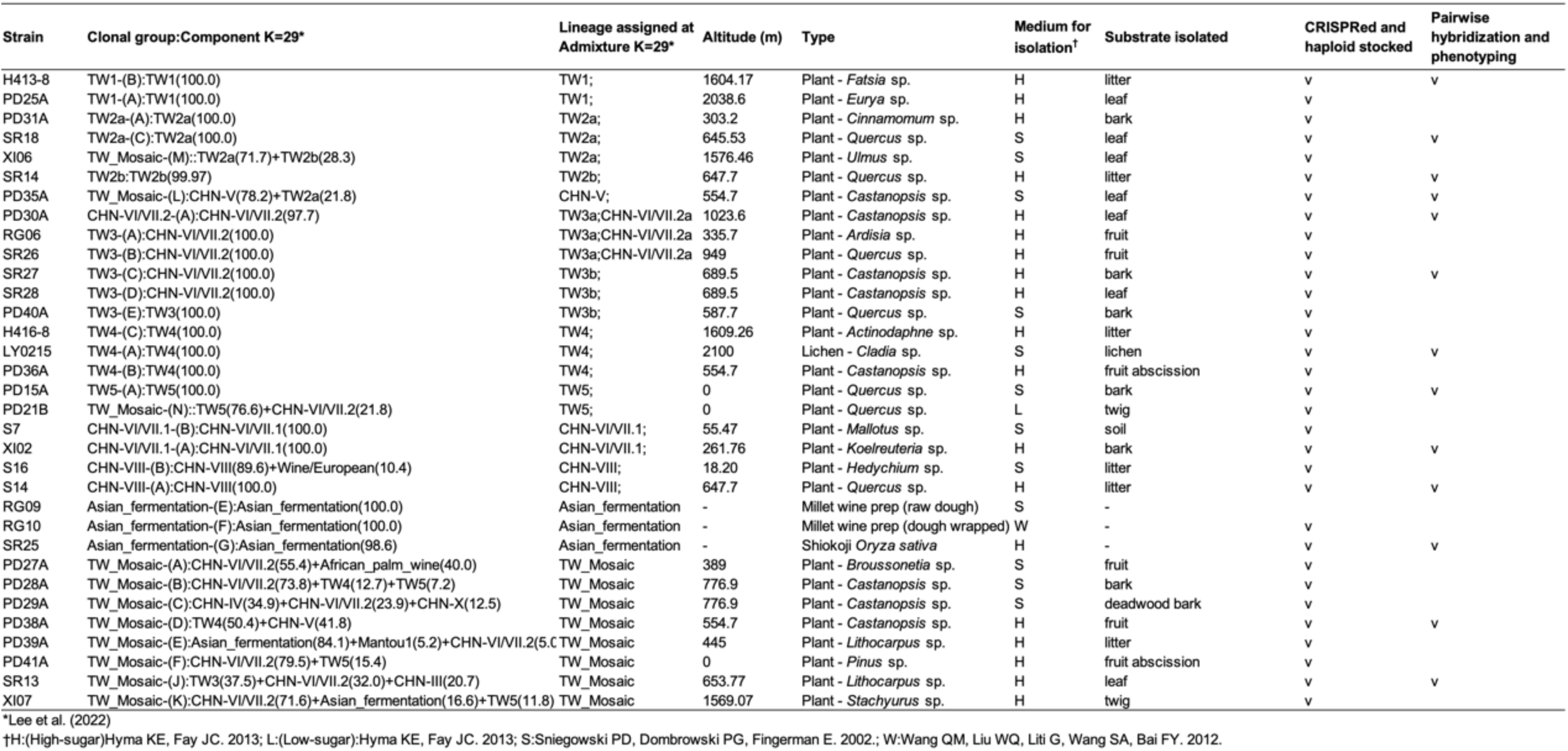
Overview of the 33 representative *S. cerevisiae* strains analyzed in this study. Stable haploids were obtained for 32 strains, of which 13 were used for pairwise hybridization and phenotyping.

Targeted disruptions of the *HO* endonuclease gene were carried out in diverse wild Taiwanese *S. cerevisiae* strains using a CRISPR-Cas9 approach (**Fig. 1c**). Positive transformants were selected using a plasmid-based kanamycin resistance marker, and successful editing was verified by PCR analysis. At least five colonies per strain were screened, all of the strains exhibited successful *HO* gene disruption, reflecting a consistently high knockout efficiency and underscoring the robustness of the CRISPR-Cas9 system in wild *S. cerevisiae* isolates.

The transformed colonies were induced to sporulate and subsequently subjected to tetrad dissection. Among the 33 strains processed with the CRISPR-Cas9 procedure, 27 were suitable for downstream genetic analyses, including 26 strains successfully produced tetrads with four viable spores, and one strain SR25 formed only dyads and triads yet still exhibited high spore viability. In contrast, five strains, including one isolate derived from fermentation products, showed markedly reduced sporulation efficiency or spore viability. Specifically, PD25A, PD31A, PD39A, RG10 and XI07 displayed spore viabilities below 50%. In addition, PD39A and the domesticated strain RG10 predominantly produced dyads or triads rather than tetrads. Another domesticated strain, RG09, failed to sporulate and exhibited pseudohyphal growth (**Supplementary Fig. S1**).

### Spore viability across Taiwanese lineages

To characterize reproductive compatibility among our wild *S. cerevisiae* isolates, we performed systematic pairwise crosses between haploids of thirteen representative strains and measured the resulting spore viability (**Fig. 2a**). Intra-strain crosses typically yielded high viability (90–100%), although SR25 was an outlier with reduced viability (59.7%), consistent with reduced sporulation efficiencies commonly observed in sake-derived strains^11,30^. Inter-strain spore viability ranged from 21% to 96%. The lowest viability was observed in crosses between SR25 (*MAT***a**) and PD35A (*MATα*), while the highest occurred between SR14 (*MAT***a**) and SR18 (*MATα*), with reciprocal crosses showing similar results.

**Figure 2.**
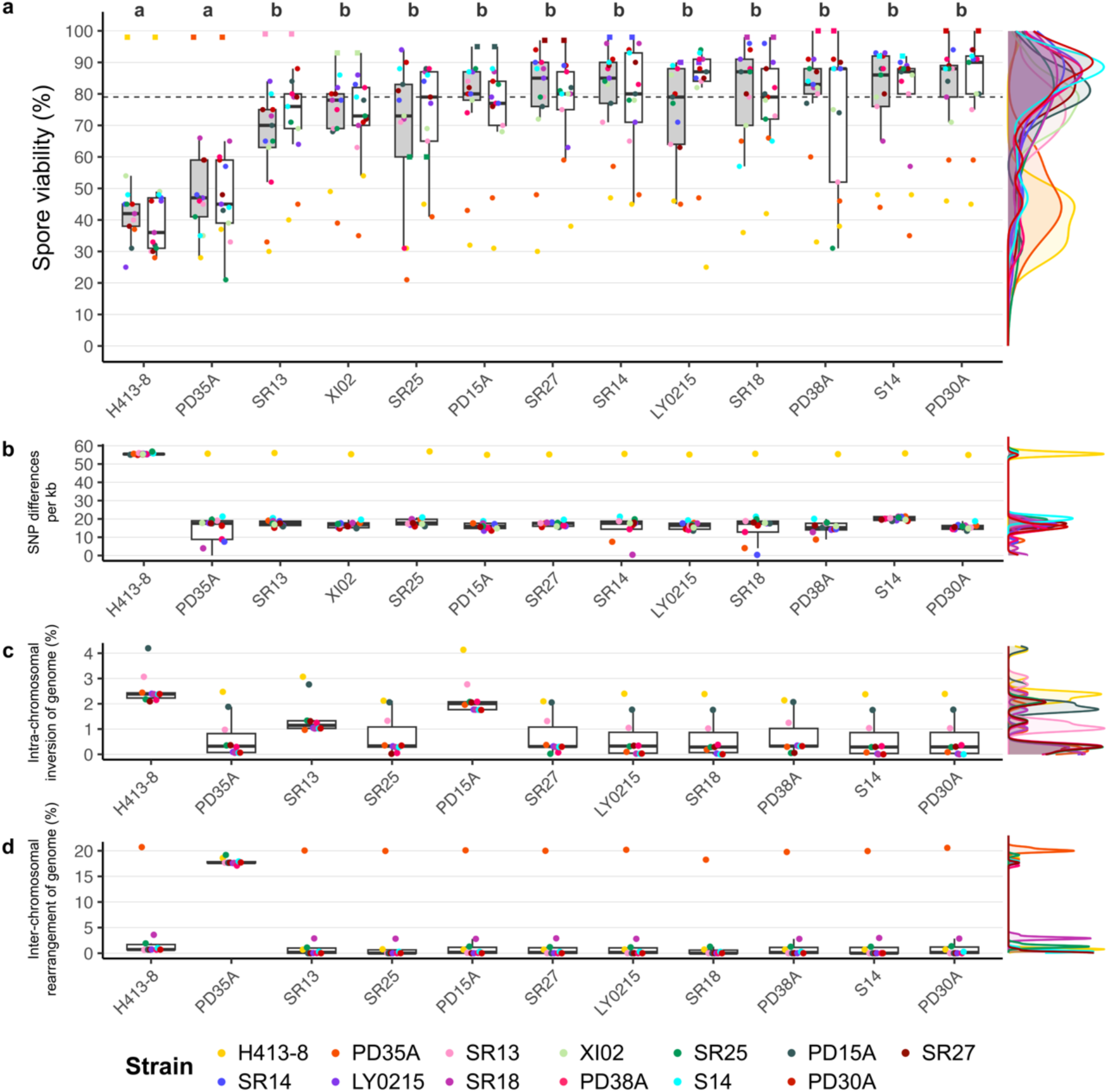
Spore viability and genomic divergence among Taiwanese *Saccharomyces cerevisiae* lineages. **a.** Spore viability (%) of pairwise crosses among thirteen representative strains. Boxplots show the distribution of spore viability across all crosses involving each strain; colored points indicate individual crosses, with colors corresponding to mating partners. Letters above boxplots indicate statistically distinct groups (ANOVA with Tukey’s HSD). The dashed line marks median spore viability. **b.** Pairwise genome-wide SNP differences between strains used in crossing experiments. **c.** Proportion of intra-chromosomal inversions between strain pairs, expressed as a percentage of the genome. **d.** Proportion of inter-chromosomal rearrangements between strain pairs, expressed as a percentage of the genome. For panels b–d, each point represents a pairwise comparison corresponding to the same strain combinations shown in panel a.

Crosses between genetically distinct lineages revealed pronounced and lineage-specific variation in spore viability. Strain H413-8, representing the most divergent Taiwanese lineage (TW1, **Fig. 2b**), consistently produced the lowest spore viability (25–49%) when crossed with all other strains (**Fig. 2a**). Similarly, PD35A (CHN-V lineage) exhibited reduced fertility (34.8–66%), despite its relatively closer genome-wide similarity to certain Taiwanese lineages such as TW2, indicating that overall sequence divergence alone does not account for reproductive isolation. Instead, these two strains were distinguished by extreme structural variation. H413-8 showed the highest proportion of intra-chromosomal inversions (mean 2.5%), followed by PD15A (2.2%), whereas PD35A exhibited a markedly elevated level of inter-chromosomal rearrangements (∼17.9%) relative to other strains (**Fig. 2c & 2d**). Consistent with this observation, spore viability was not significantly correlated with genome-wide genetic distance once these outliers were considered (**Supplementary Fig. S2**), indicating that reproductive isolation in Taiwanese *S. cerevisiae* is driven by a limited number of major genomic incompatibilities rather than gradual divergence. Such barriers are in line with previous studies demonstrating that chromosomal rearrangements and locus-specific incompatibilities can severely disrupt meiosis even among geographically proximate yeast populations^25,31–33^. Together, these patterns suggest that many wild *S. cerevisiae* lineages persist primarily through asexual reproduction, with sexual cycles occurring infrequently and contributing little to ongoing gene flow despite spatial overlap^14,34^.

In contrast to these strongly isolated lineages, natural hybrids exhibited comparatively high fertility. The natural hybrid strain SR13 demonstrated hybrid genomic features (TW3 (37.5) + CHN-VI/VII.2 (32.0) + CHN-III (20.7)) with moderate to high spore viability (52–88%), further highlighting the ecological and evolutionary relevance of hybridization among wild yeast lineages. Examination of mating-type combinations uncovered intriguing asymmetries in viability between *MATα*-*MAT***a** and *MAT***a**-*MATα* crosses (**Fig. 2a**). Although most crosses exhibited similar viabilities irrespective of mating direction, a few exceptions were observed. The natural hybrid PD38A (TW-mosaic) exhibited pronounced mating-type effects: when acting as the *MATα* parent, spore viability varied widely across partners (31–91%), whereas when serving as the *MAT***a** parent, all crosses consistently showed high viability (>77%). These asymmetric patterns indicate mating-type-specific interactions influencing hybrid fertility, potentially mediated by genetic or epigenetic differences associated with the *MAT* locus or its regulatory context.

### Thermal phenotypic variation of wild *S. cerevisiae* and their natural hybrids in Taiwan

Building on the observed reproductive and lineage-specific differences, we performed growth assays across a temperature gradient (**Fig. 3**). Most strains exhibited similar optimal growth temperatures, consistent with their shared subtropical environments^14,35^. However, distinct lineage-specific patterns emerged. Strain H413-8 from lineage TW1 consistently displayed slightly lower optimal growth temperatures (31.4 ° C) compared to other lineages (32.3°C–37.1°C), placing it closer to the thermal optima observed in other species of the *Saccharomyces* genus, such as *S. paradoxus*, and suggesting retention of a more ancestral thermal trait^36^. Interestingly, natural hybrid isolate PD38A demonstrated notably higher optimal growth temperature of 35.1°C, compared to the strains LY0215 (34.1°C), PD35A (34°C) that belong to its parental lineages (TW4, CHN-V respectively) and wider TPB_80_ (defined as the temperature range over which performance remains at or above 80% of the maximum growth rate; PD38A, 13.7°C; LY0215, 12.5°C; PD35A, 12.1°C). SR25 (Asian fermentation), isolated from Shiokoji fermentation practice, had the highest T_opt_ of 37.1°C and highest μ_max_ of 0.32 h^−1^, on the other hand, it had the most limited TPB_80_ (10.4°C) among all the tested strains.

**Figure 3.**
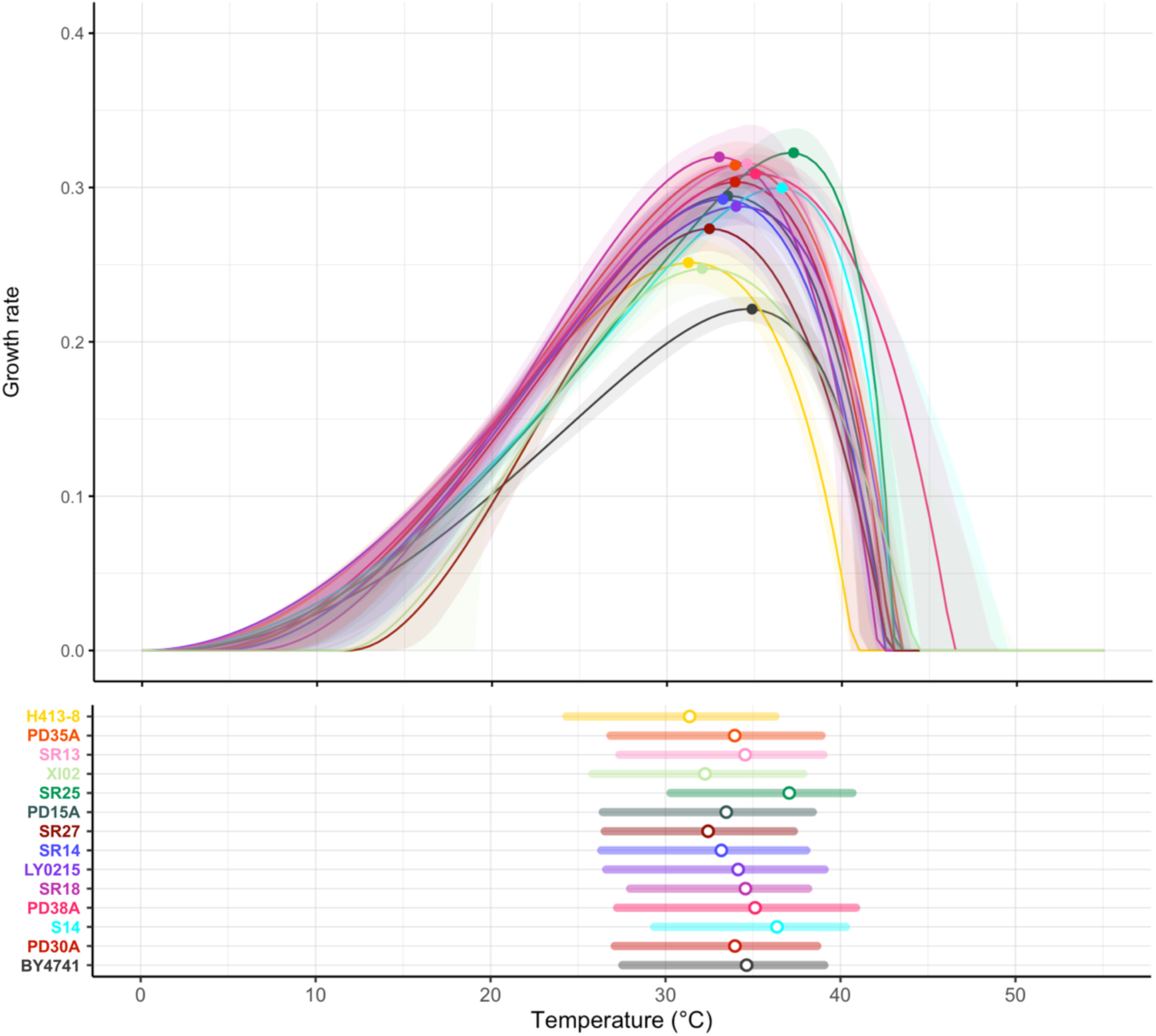
Thermal performance curve (TPC) and thermal performance breadth 80% (TPB_80_) of 13 representative strains and BY4741.

Across 11 non-anthropogenic strains (excluding SR25 and S14), optimal growth temperature was significantly positively correlated with maximum growth rate (R² = 0.58, P < 0.01; **Supplementary Fig. S3**), consistent with a “hotter is better” pattern^36^. However, no significant relationship was detected between optimal growth temperature and thermal performance breadth (TPB₈₀) (**Supplementary Fig. S4**), indicating that higher thermal optima did not correspond to broader performance ranges. Thus, within wild Taiwanese *S. cerevisiae*, the “hotter is wider” hypothesis was not supported^36^.

### Growth curve under different carbon sources

We assessed the growth of 13 representative strains and BY4741 under a variety of media containing different carbon sources at 30°C. Among six different carbon source media, the growth in maltose had the lowest median μ_max_ (**Fig. 4a**). All 13 representative strains had significantly higher μ_max_ compared to lab strain BY4741 when growing under glucose (F_13,61_ = 9.69) and raffinose (F_13,61_ = 92.93) based medium (**Supplementary Table S1**). SR27 (TW3b) had the highest μ_max_ when in fructose based medium, and it was significantly higher than all the others (F_13,61_ = 15.95). On the other hand, SR18 (TW2a) had the highest μ_max_ in three carbon sources, which is glucose, raffinose, galactose under 30°C. H413-8 (TW1) had the lowest μ_max_ within 13 representative strains when in fructose, glucose, sucrose, and maltose, and the latter even lower than lab strain BY4741. In the assessment of maltose utilization, S14 (CHN-VIII) and SR25 (Asian fermentation) exhibited significantly higher maximum growth rates (μ_max_) of 0.18 and 0.12 h^−1^ (F_13,61_ = 340.5), with S14 being significantly higher than the domesticated strain SR25 (P < 0.001). Principal component analysis based on μ_max_ values across all carbon sources (**Fig. 4b**). The first two components explained 67.4% of the variance (PC1 = 44.4%, PC2 = 23%). Strains with enhanced maltose utilization, including S14, SR25, and SR27, clustered toward one end of the PCA space, whereas H413-8 and XI02 grouped separately, reflecting their generally reduced growth across most substrates. Together, these results demonstrate pronounced metabolic differentiation among wild *S. cerevisiae* lineages.

**Figure 4.**
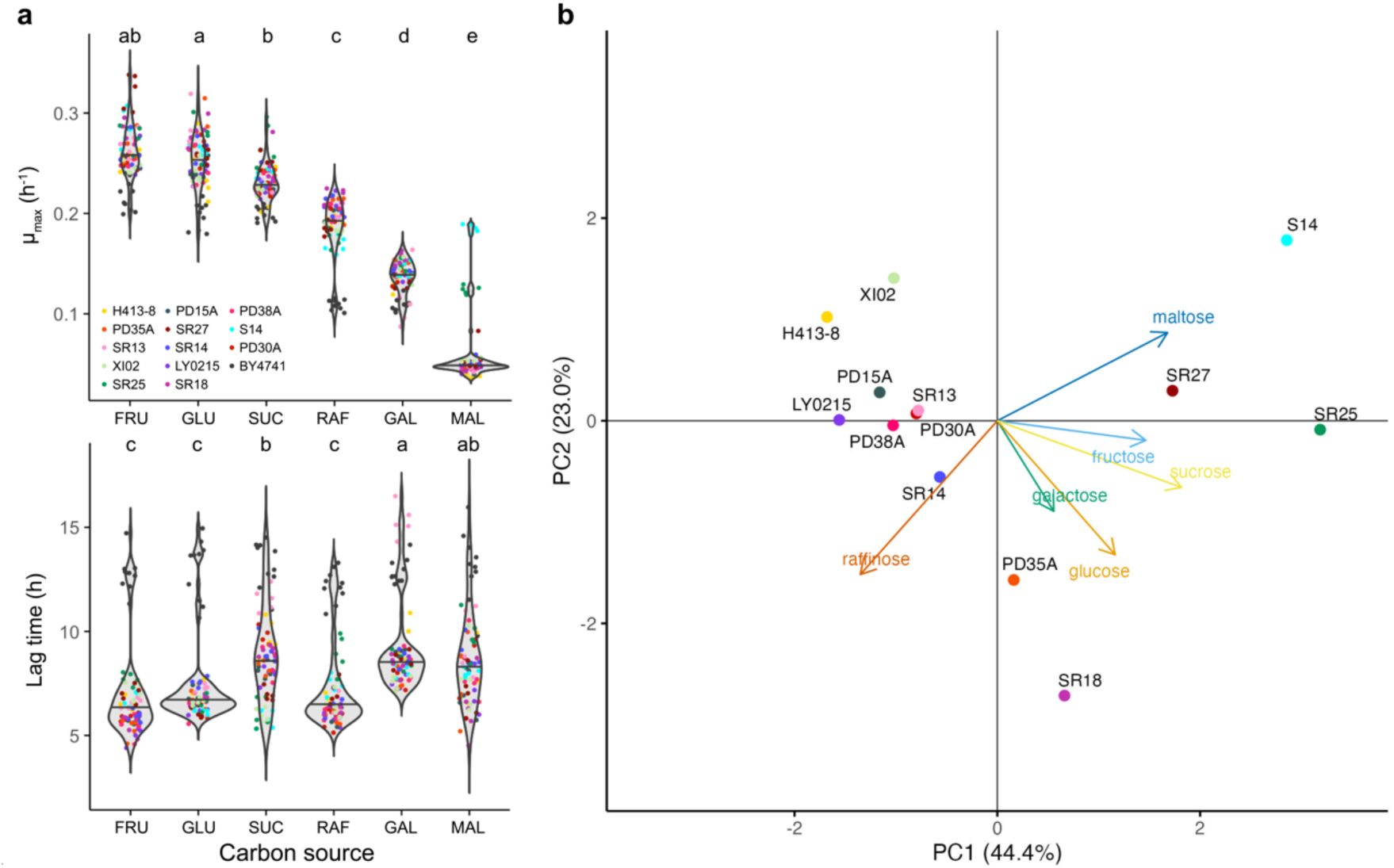
Characteristics of yeast strain growth under different carbon sources. **a.** Growth metrics of representative strains and BY4741. μ_max_ was calculated from the growth curve up to the stationary phase; lag time was conducted within 24 h yeast growth. GLU = glucose; FRU = fructose; SUC = sucrose; RAF = raffinose; GAL = galactose; MAL = maltose. Letters above boxplots indicate statistically distinct groups (ANOVA with Tukey’s HSD). **b.** Principal component analysis of representative strains, using maximum growth rate under different carbon source. Arrows indicate different carbon source in the media.

### Genomic basis of maltose utilization in S14 and SR25

Given the distinct maltose catabolization profiles of S14 and SR25, both associated with anthropogenic or feral contexts, we examined genomic variation at the *MAL* loci using available long-read assemblies from Taiwanese strains^14^. These loci are typically located in the highly dynamic subtelomeric regions of chromosome VII. Synteny analysis suggested that the ancestral configuration of the chromosome VII *MAL* region likely comprised two copies of *IMA2*, four *MALR* homologs, three *MALT* genes, and a single *MALS* gene (**Fig. 5a**). SR25 carried three complete *MAL* loci located at the subtelomeric ends of chromosomes VII, II, and VIII, each retaining intact *MALR–MALT–MALS* clusters. In contrast, most other Taiwanese strains and the laboratory strain BY4741 possessed only an additional *MAL* locus on chromosome II, in which *MALT* and *MALS* were frequently truncated or absent (**Supplementary Fig. S5**). These patterns are consistent with extensive subtelomeric restructuring. For example, PD35A, which exhibits substantial chromosomal rearrangements, lacked the homologous *MAL13–AGT1–IMA2* region entirely. Nevertheless, comparable *MAL* copy numbers were observed among strains (seven copies in S14, sixteen in SR25 and six to nine in the remaining strains), suggesting that copy number alone is insufficient to explain the observed differences in maltose growth.

**Figure 5.**
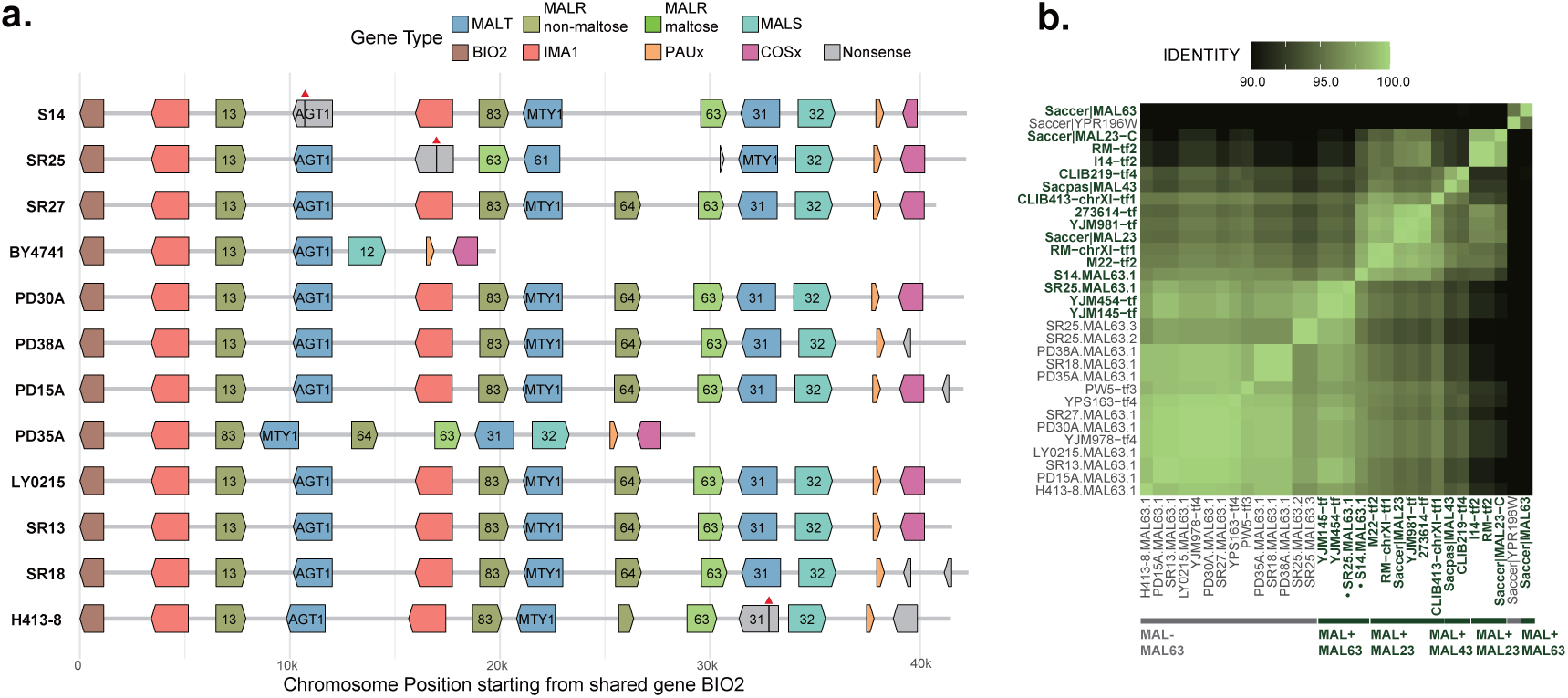
*MAL* loci divergence in *Saccharomyces cerevisiae* strains lead to differences in maltose proficiency. **a.** Synteny of the *MAL* loci on chromosome VII. Genes are colored by gene families and *MAL* genes are labeled by homology group. **b.** Amino acid sequence identity heatmap of *MAL63* homologs. *MAL*^+^ *MALR*s are labeled with dark green color.

Previous studies have shown that *MAL* homologs differ in their functional contribution to maltose and other carbon source utilization^37–39^, with allelic variation in the transcriptional regulator *MAL63* playing a central role in determining maltose proficiency. Consistent with this, multiple sequence alignment revealed that *MAL63* alleles from S14 and SR25 clustered with previously characterized *MAL*⁺ variants and showed high sequence similarity to known functional alleles (**Fig. 5b; Supplementary Fig. S6**). In both strains, maltose-specific *MALT* genes (**Fig 5a; Supplementary Fig. S7**) were also located in close genomic proximity to *MAL*⁺ *MAL63* alleles on chromosome VII. Together, these results indicate that enhanced maltose utilization in S14 and SR25 is associated with functional *MAL*⁺ regulatory alleles and linked maltose-specific transporters. Interestingly, SR27, which exhibited the third-highest maltose growth rate, carried a *MAL63* allele that grouped with *MAL*⁻ variants, suggesting that SR27 may utilize non-canonical regulatory pathways or unidentified transporters to achieve maltose proficiency.

## Discussion

Our haploid collection of wild Taiwanese *S. cerevisiae* enabled direct assessment of reproductive compatibility and phenotypic diversity across genetically and ecologically distinct lineages, even though these strains are often sympatric. This systematic comparison revealed striking variation in hybrid fertility, even among strains that coexist in the same habitats. Spore viability across inter-lineage crosses ranged from 21% to 96%, with a mean of 69.8%, broadly consistent with the range previously reported in a review by Bendixsen et al.^40^, which documented values from 2.5% to 100% with a mean of 75%. The most divergent lineages—H413-8 (TW1) consistently produced low-viability hybrids, indicating strong lineage-specific reproductive barriers.

We had previously observed that wild *S. cerevisiae* lineages in Taiwan reproduce predominantly asexually^14^, and our current findings now clarify one of the key reasons behind this pattern. In all hybrid combinations tested, zygotes exhibiting the characteristic trilobed morphology were observed, indicating that mating and cell fusion occurred successfully. This suggests that prezygotic isolation is minimal or absent, and that reproductive barriers in these lineages primarily act at the postzygotic stage. This observation aligns with previous work showing that postzygotic mechanisms, such as reduced spore viability, are the predominant form of reproductive isolation in *Saccharomyces*^33^. Consistent with earlier studies, structural rearrangements and locus-specific incompatibilities represent major causes of postzygotic isolation in yeasts^25,31–33^, highlighting the role of genomic structural variation in maintaining the separation of these lineages despite geographic coexistence. Having identified the genetic constraints that favor asexual reproduction, the next question is whether environmental conditions in these subtropical forests further encourage this lifestyle by selecting for robust mitotic growth and local adaptation over recombination and dispersal.

Natural hybrids within our collection exhibited high spore viability across various strains, exemplified by strain PD38A, a hybrid between TW4 and CHN-V lineages. Such hybrids may benefit from beneficial recombination events or alleviated genetic incompatibilities, conferring adaptive advantages under fluctuating or stressful conditions^40–42^. The fertility and stability of PD38A underscore the evolutionary role of hybridization in wild *S. cerevisiae*, where occasional gene flow may introduce beneficial variation and enhance ecological resilience among otherwise isolated lineages. Intriguingly, PD38A also revealed asymmetric fertility between reciprocal crosses: when acting as a *MATα* parent, its hybrids showed markedly lower viability than when serving as *MAT***a**. Such directional effects suggest mating-type-specific interactions influencing reproductive compatibility, potentially mediated by genetic or epigenetic variation near the *MAT* locus^43,44^. These results highlight that while most wild lineages remain reproductively isolated, rare hybridization events—shaped by both genomic and mating-type constraints—can serve as key catalysts of genetic exchange in natural yeast populations.

Phenotypically, our isolates exhibited marked variation in thermotolerance, a trait critical for ecological adaptation and evolutionary fitness^36,45^. Although most strains shared similar optimal growth temperatures consistent with subtropical climates, lineage-specific differences were evident. The TW1 strain H413-8 showed a lower optimal temperature that overlapped with other *Saccharomyces* species, suggesting retention of an ancestral thermal phenotype^36,45–48^. In contrast, the natural hybrid PD38A displayed elevated thermotolerance and broader thermal performance breadth relative to its parental lineages, consistent with hybridization facilitating ecological expansion^27,49^.

Across wild Taiwanese strains, we observed a clear “hotter is better” pattern, with higher optimal temperatures associated with increased maximum growth rates, mirroring interspecific trends across the *Saccharomyces* genus^36^. However, we found no evidence for a “hotter is wider” relationship within *S. cerevisiae*, as higher optima did not correspond to broader thermal breadth. Riles & Fay^49^ demonstrated that polymorphisms in stress-responsive genes contribute to thermotolerance variation in wild *S. cerevisiae*, providing a genomic basis for such performance differences. Together, these results suggest that thermal adaptation within species primarily enhances peak performance rather than expanding ecological tolerance, indicating that niche breadth may remain constrained by lineage-specific genetic and physiological trade-offs.

Variation in carbon source utilization among wild lineages further highlights the ecological diversification of *S. cerevisiae*. Most wild strains grew efficiently on simple sugars such as glucose and fructose but exhibited poor growth on maltose, consistent with the limited maltose metabolism typically observed in non-domesticated yeasts^8,45^. Genomic analyses revealed that enhanced maltose utilization in the outliers S14 and SR25 was associated with functional *MAL*⁺ regulatory alleles and linked maltose-specific transporters, whereas SR27 achieved relatively high maltose growth despite carrying a *MAL63* allele that grouped with *MAL*⁻ variants, indicating that alternative genetic routes can also support maltose utilization. These findings refine and extend previous observations that *MAL* gene copy number alone does not predict maltose proficiency^37,38^, showing that both canonical *MAL*⁺ regulatory alleles and alternative genetic routes can underlie enhanced maltose utilization in wild and feral *S. cerevisiae* lineages. Integrating phenotypic assays with long-read-resolved genomes links ecological performance to specific loci and alleles, revealing a mosaic of local adaptations in which feral and hybrid strains retain broader metabolic flexibility than most wild lineages^12^.

In summary, the haploid strain collection developed in this study provides a robust genetic toolkit for exploring the ecological and evolutionary dynamics of wild *S. cerevisiae*. By enabling direct analyses of reproductive compatibility, hybridization, and phenotypic diversity across natural lineages, this resource bridges the gap between classical yeast genetics and population-level ecology. Our results reveal how reproductive barriers, hybrid fertility, and lineage-specific adaptations collectively shape the diversification of wild yeast populations. This also highlighted the intraspecies differences. Integrating these genomic insights with ecological context highlights how both intrinsic incompatibilities and environmental selection contribute to the maintenance of a predominantly asexual lifestyle in nature. This framework opens new avenues uniting classical genetics with natural variation to uncover how microevolutionary processes (like hybridization and local adaptation) generate biodiversity. The haploid wild yeast resource we developed will facilitate further research into the genetic and ecological mechanisms that drive evolution in microbes, offering a foundation for both fundamental discovery and biotechnological innovation.

## Materials and Methods

### Strain selection and stable haploids construction

Yeast sampling and enrichment were described in Lee et al^14^. In brief, we collected 2,461 environmental samples from various substrates. We chose 33 non-clonal wild yeast strains that were representative of the yeast genetic diversity (**Table 1**). All strains were maintained on YPD agar (1% yeast extract, 2% peptone, 2% glucose (dextrose), 2% agar) and were preserved in 25% glycerol at -80°C.

Stable haploids were generated from wild *S. cerevisiae* strains, which are predominantly homozygous diploids (averaging heterozygosity of 2.57 × 10^−4^) and homothallic, by targeted *HO* gene knockout using CRISPR/Cas9 approach^28,29^. Briefly, plasmid pL88 carrying a kanamycin resistance gene was used to introduce two cuts at the *HO* locus. Homology-directed repair templates for the *HO* locus were generated by PCR using oligos HO_truncated_F (5’-TTT AAA ATG CTT TCT GAA AAC ACG ACT ATT CTG ATG GCT AAC GGT GAA ATA TCG AGT ATG TGC TAG ATG CTA TGG AAG ATA CAA ATT CAG CGG TCA TCA C-3’) and HO_truncated_R (5’- GTG ATG ACC GCT GAA TTT GTA TCT TCC ATA GCA TCT AGC ACA TAC TCG ATA TTT CAC CGT TAG CCA TCA GAA TAG TCG TGT TTT CAG AAA GCA TTT TAA A-3’). After the transformation, at least five positive colonies were preserved and checked for *HO* genotype. To confirm the targeted *ho* locus had been successfully edited, PCR was performed using primer pair HO_165F (5’-GTT GAA GCA TGA TGA AGC G-3’)/HO_1702R (5’-CAA ACT GTA AGA TTC CGC CAC-3’). Size of the amplicon from successful colonies should be 398 bp while the original colonies yielded 1867 bp. One successfully-edited colony from each strain was further subjected to sporulation and tetrad dissection to acquire haploid strains. Sporulation of *ho*-knockout diploids was performed induced on a solid sporulation medium, composed of 1% potassium acetate, 0.1% yeast extract, 0.05% glucose, 0.01% dropout powder^50^ and 2% agar, incubated for 3–7 days. Mature asci were then digested with zymolyase (Zymo Research, California, USA) to digest the ascus wall, and tetrad dissection was performed using Singer MSM system series 300. Only tetrads in which all four spores were viable were selected for further analyses, except for strain SR25, which failed to form tetrads.

### Mating type analysis

Mating types of spores were determined using *S. cerevisiae* tester strain *bar1*Δ and *sst2*Δ. The *bar1*Δ strain is sensitive to α-factor, while the *sst2*Δ strain is sensitive to a-factor. Single colonies of each tester were resuspended in 200 μL ddH_2_O and spread on YPD agar supplemented with 0.01% Tryptophan, 0.01% Adenine. For the *sst2*Δ tester, the medium additionally contained 0.04% Triton X-100 to enhance halo formation. Colonies from each spore were resuspended in 5 μL ddH_2_O and 2 μL of the mixture was spotted onto the *bar1*Δ and *sst2*Δ tester lawns, which were then incubated for 2 days at 30°C. Mating type was inferred from the presence of a growth inhibition halo on the corresponding tester lawn. Only tetrads producing two *MAT***a** and two *MATα* spores were retained and haploids were preserved in 25% glycerol at -80°C for subsequent analyses, except for strain SR25, which failed to form tetrads.

### Genome sequencing

*S. cerevisiae* strains were cultured in 30°C YPD broth for 1 day with 200 rpm agitation before gDNA extraction, then pelleted by centrifuge. For Illumina sequencing, gDNA was extracted with Zymo Quick-DNA Fungal/Bacterial Miniprep Kit (Zymo Research, California, USA) following the manufacturer’s instruction. Library preparation and sequencing were outsourced by Biotools Co. and performed on Novaseq 6000 platform (**Supplementary Table S2**).

For Oxford Nanopore Technologies (ONT) sequencing, high-molecular-weight gDNA was extracted. Briefly, yeast cell wall was digested with zymolyase for spheroplasting, followed by cell lysis using lysis buffer supplemented with sodium dodecyl sulfate (SDS). Proteins were subsequently precipitated, and gDNA was recovered by DNA precipitation. ONT library was prepared via ligation sequencing kit (SQK-LSK109) and native barcoding (EXP-NPD104). Nanopore reads were then sequenced on an R10.3 flow cell (FLO-MIN111) and base-called using Guppy (v5.0.22). Assembly method and assemblies for strains S14, LY0215, PD15A, PD35A, PD38A, SR18, PD35A were derived from a previously published study^14^ (**Supplementary Table S3**).

### Variant analysis

Raw 150 base-pair paired-end Illumina reads for each individual were trimmed with fastp v0.24.0^51^ and aligned to reference *S. cerevisiae* genome S288C (GenBank accession: GCA_000146045.2) using the default parameters of BWA-MEM2 v2.2.1^52^. Duplicated reads were marked in the sorted alignment map files using samtools v1.1.19^53^ and GATK MarkDuplicates v4.4.0.0^54^. Single-nucleotide polymorphisms (SNPs) were detected using bcftools v1.1.19^53^ and variant loci with more than 10% missing genotypes were filtered. Genetic distance and pair-wise SNP difference on 10-kb windows were calculated using pixy v1.2.11^55^. To identify structural variants (SVs), including chromosomal inversions and large-scale rearrangements, we performed pairwise whole-genome alignments among *S. cerevisiae* strains for which long-read assemblies were available^14^. Strains XI02 and SR14 were excluded from this analysis due to the absence of long-read sequencing data. Genome alignments were generated using nucmer from the MUMmer v3.23 package^56^ with default parameters. The resulting delta alignments were filtered using delta-filter (–u 50, –l 300) to retain unique alignments and segments of at least 300 bp. Filtered alignment coordinates were extracted using show-coords (–cdHlrT) and subsequently analyzed in R.

### Yeast crosses and spore viability assays

Pairwise intraspecific crosses were performed among selected *MAT***a** and *MATα* haploid strains to assess reproductive compatibility. Single colonies of each strain were streaked on YPD and incubated at 30°C overnight. Haploid cells from each pair were mixed in 5 µL of sterile ddH_2_O, and the suspension was then transferred onto a YPD plate and incubated at 30°C for 3 hrs to allow mating. Diploid zygotes were identified and isolated using a Singer MSM 300 micromanipulator based on their characteristic trilobed budding morphology (**Fig. 1c**), indicative of the first post-zygotic division. Diploid colonies were incubated at 30°C for 2 days and transferred to 1% potassium acetate agar to induce sporulation. After sporulation, tetrads were dissected as described above using the Singer MSM 300 system. For each cross, at least 28 tetrads (112 potential spores) were dissected, and spore viability was calculated as the percentage of spores that germinated and formed visible colonies on YPD. Permutation tests assessing the relationships between spore viability, sequence divergence, and genomic structural variation were performed using the Freedman–Lane method as implemented in the R package permuco, with 10,000 permutations.

### Quantitative growth phenotyping

Mitotic growth was evaluated under varying temperatures and carbon sources. Temperature-dependent growth was measured in liquid YPD (1% yeast extract, 2% peptone, 2% dextrose) across a gradient of 20, 25, 30, 32.5, 35, 37.5, 40, and 42°C. Carbon source utilization was assessed in media containing 1% yeast extract, 2% peptone, and 2% of a single carbon source (glucose, fructose, galactose, sucrose, maltose, or raffinose). All media were filter-sterilized, and pH values ranged from 5.8 to 6.0.

Each strain was pre-cultured in 100 µL YPD at 25°C for 24 h with agitation to saturation, then diluted 1:10,000 into 96-well U-bottom microplates (Corning Costar, tissue-culture treated) to a final volume of 150 µL (∼1.5 × 10³ cells per well). To avoid edge effects from evaporation, edge wells were not used for strain inoculation, but were filled with the same medium and excluded from analysis to minimize evaporation effects. Strain positions were randomized within each plate. Growth (OD_595_) was monitored every 10 min using a Tecan Infinite M200 Pro microplate reader until stationary phase. Each condition was tested in five independent replicates derived from separate colonies.

Growth curves were smoothed using the gcplyr R package^57^ with a moving-median function. Maximum growth rate (μ_max_), lag time and other metrics were estimated using model-free analysis in R4.4.1. Thermal performance curves (TPCs) were fitted using the Cardinal Temperature Model with Inflection (CTMI) as described by Molinet & Stelkens^36^, starting from T_min = 0 °C, T_opt = 31 °C, T_max = 42 °C, and R_opt = 0.25. From each fitted model, the optimal growth temperature (T_opt_), minimal growth temperature (T_min_), maximum growth temperature (T_max_), maximum performance, and thermal performance breadth (the temperature interval across which performance exceeds 80% of μ_max_, TPB₈₀) were derived. Confidence intervals were estimated by 1,000 bootstrap replicates.

### Genomic annotation and comparative analysis of *MAL* loci

To determine the genomic factors affecting maltose growth, available genomes generated by ONT long reads from Lee et al.^14^ were annotated using the funannotate2 v25.11.1^58^ pipeline trained on *Saccharomyces cerevisiae* S288C annotation SGD R64-4-1. Homology groups of funannotate2-curated *MAL* genes were assigned through multiple sequence alignment with reference and published gene sequences^38^ using MAFFT^59^. Pairwise amino acid sequence identity was calculated by BLAST+^60^. GFFs covering the homologous subtelomeric *MAL* regions were transformed to BED files using AGAT v1.4.2^61^ and plotted on R v4.3.2.

## Acknowledgement

We thank Dr. Jun-Yi Leu for providing the tester strains. IJT is funded by National Science and Technology Council, Taiwan (Grant NSTC 113-2628-B-001-002 and 114-2628-B-001-014) and Academia Sinica (Grant AS-IA-113-L04).

## Authors’ contributions

IJT and HMK conceived the study. GL designed the CRISPR–Cas9 constructs. CJY and CH conducted the majority of the experiments, including gene engineering, yeast mating, phenotyping, and manuscript drafting. YCY analyzed the genome sequences. YHL performed part of the experiments related to gene engineering and yeast mating. CJY and MRL analyzed the thermal performance datasets. IJT, HMK and CJY wrote the manuscript with inputs from all authors.

## Data Availability

**Supplementary Table S2**. **and S3** summarize the sequencing data presented in this study. The data can also be accessed under accession PRJNA1420517, or PRJNA755173 for published Nanopore assemblies.

## Supplementary Information

**Supplementary Figure S1.**
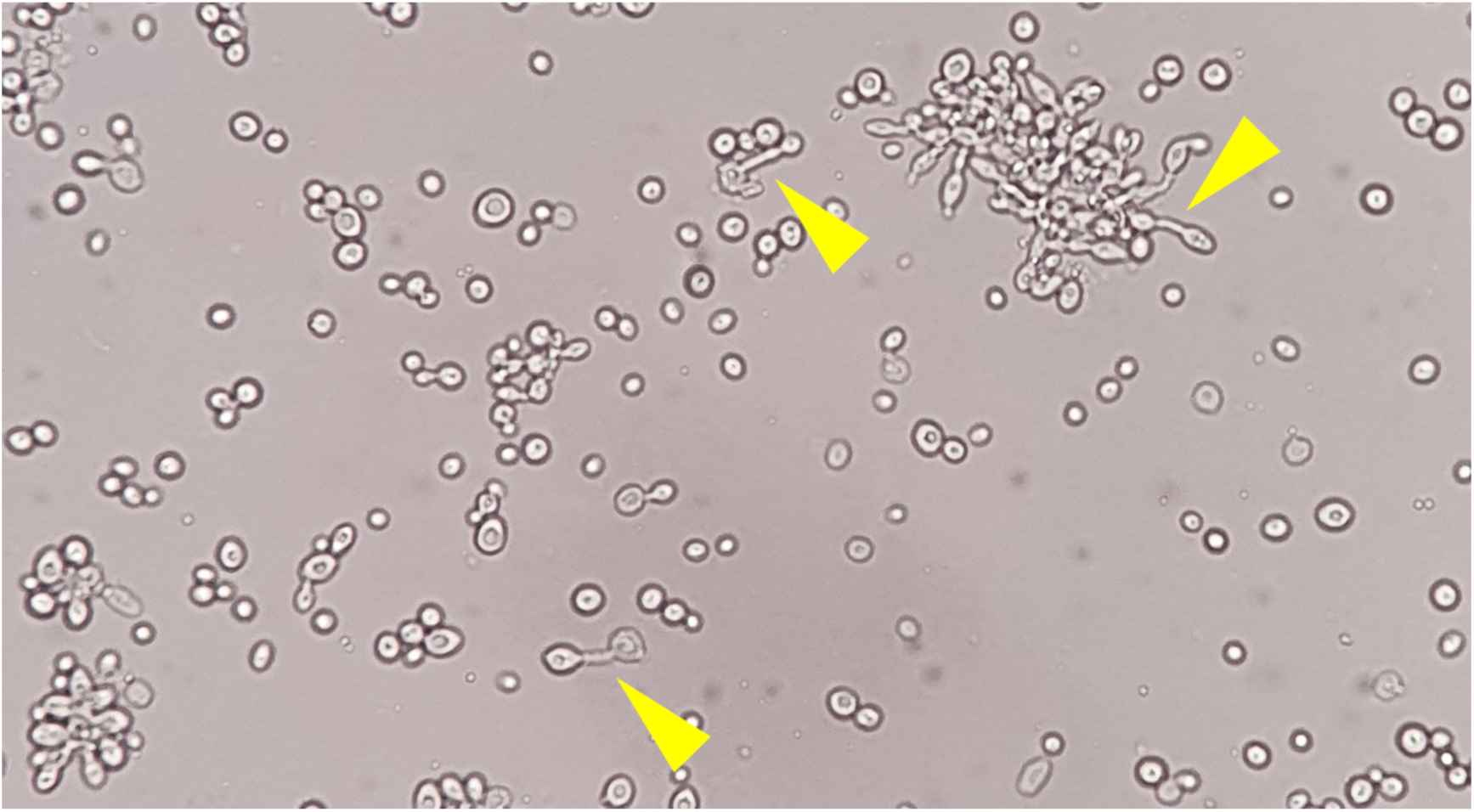
Pseudohyphal growth of *HO* knockouted RG09.

**Supplementary Figure S2.**
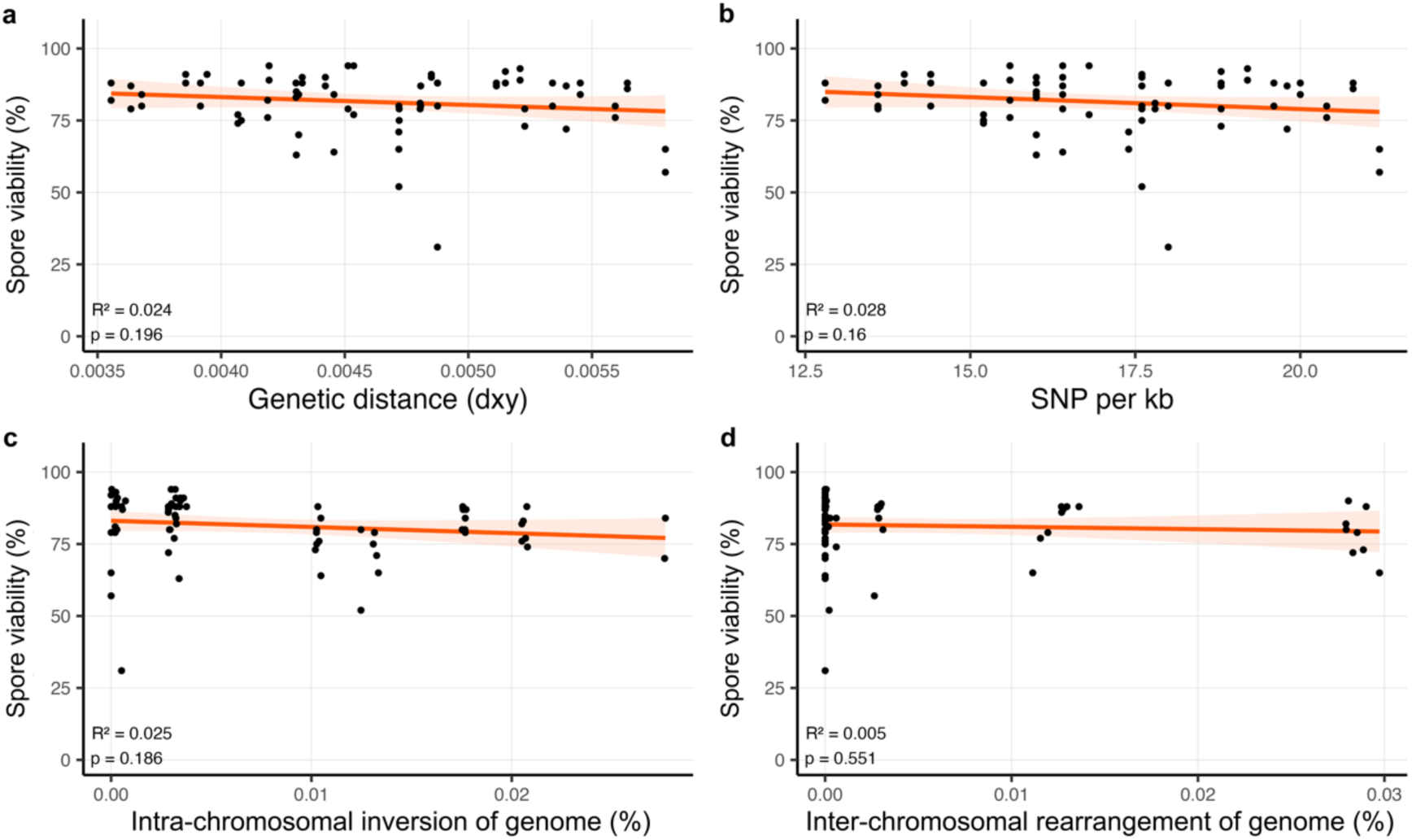
Regression of spore viability in pairwise hybridizations against different metrics. **a.** genetic distance (dxy)**; b.** SNP differences per kb**; c.** intra-chromosomal inversion between strains; **d.** inter-chromosomal rearrangement between strains. Each facet is labeled with *MAT***a** parental strains. Self-crossing and outliers were not included.

**Supplementary Figure S3.**
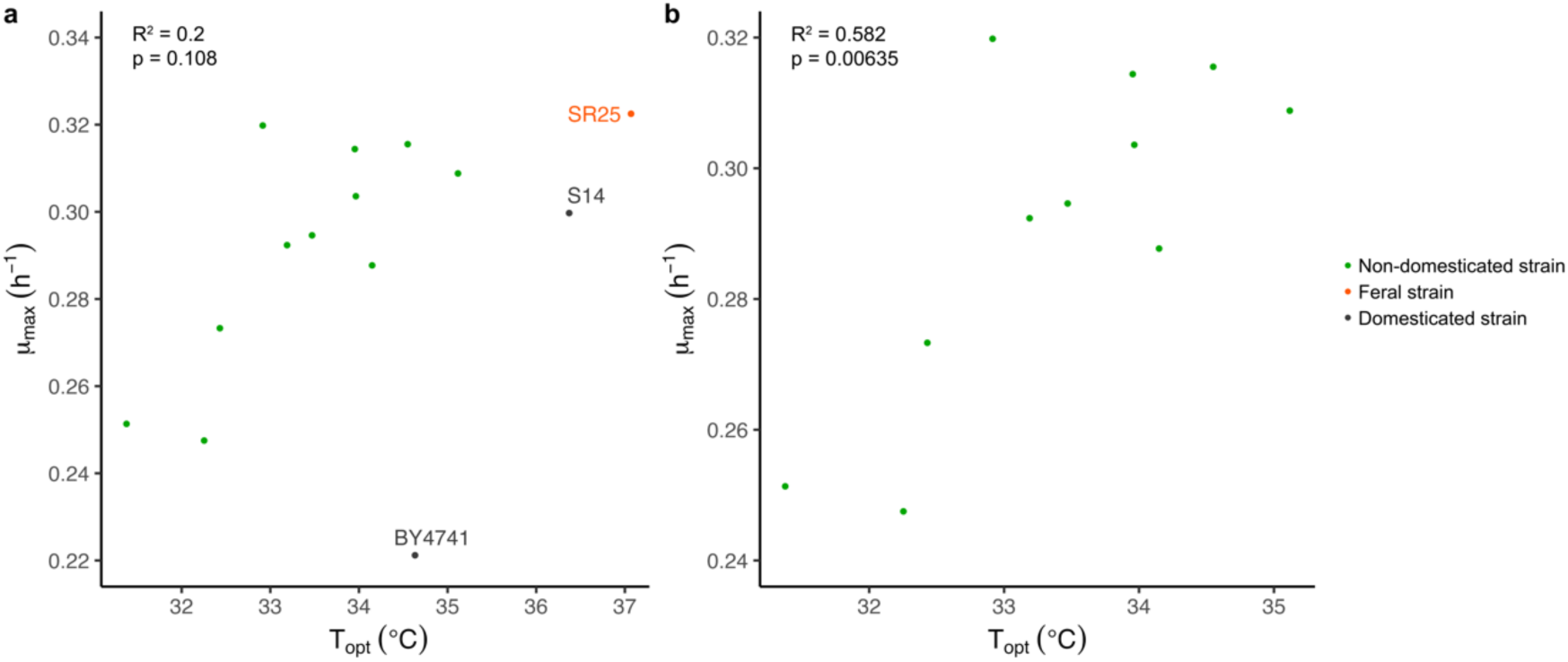
Correlation between optimum temperature (T_opt_) and maximum growth rate (μ_max_). **a.** Correlation of 13 representative strains and lab strain BY4741; **b.** 11 representative predomesticated yeast strains.

**Supplementary Figure S4.**
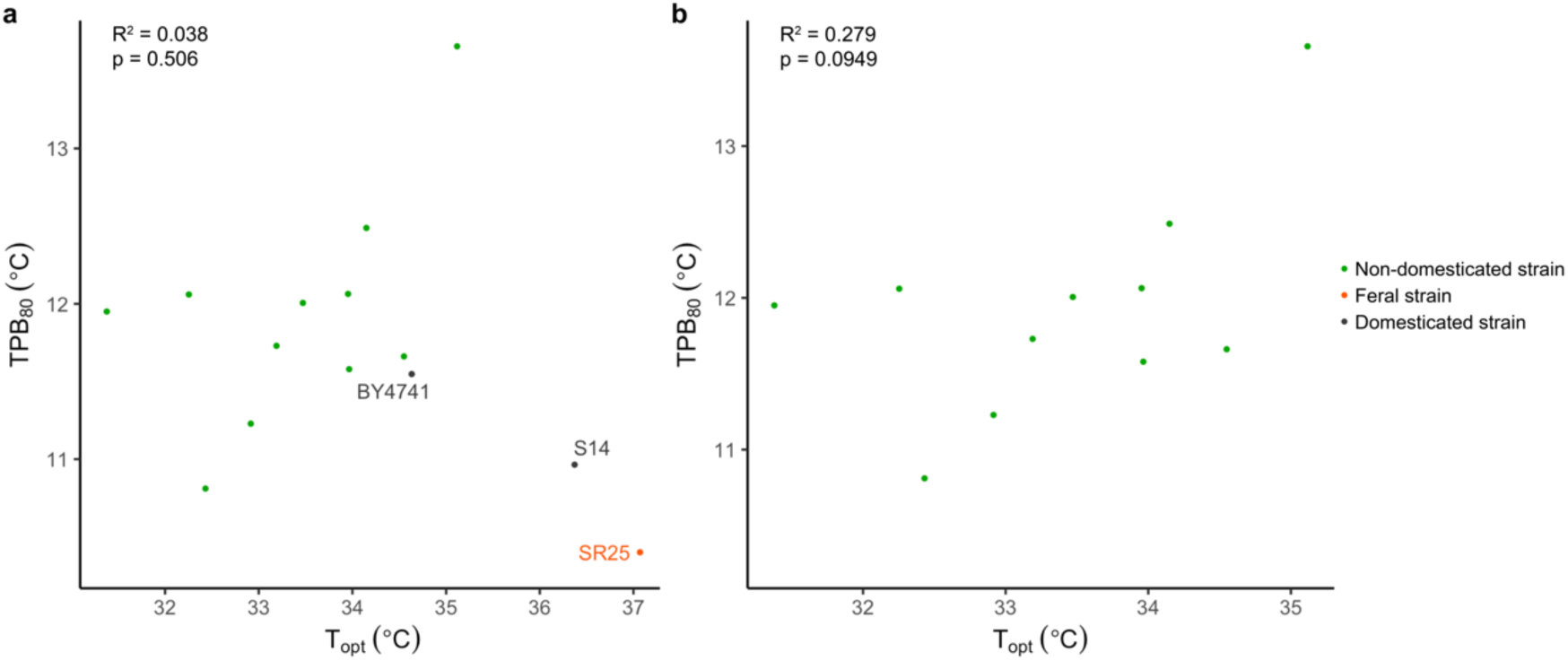
Correlation between optimum temperature (T_opt_) and 80% thermal performance breadth (TPB_80_). **a.** Correlation of 13 representative strains and lab strain BY4741; **b.** 11 representative predomesticated yeast strains.

**Supplementary Figure S5.**
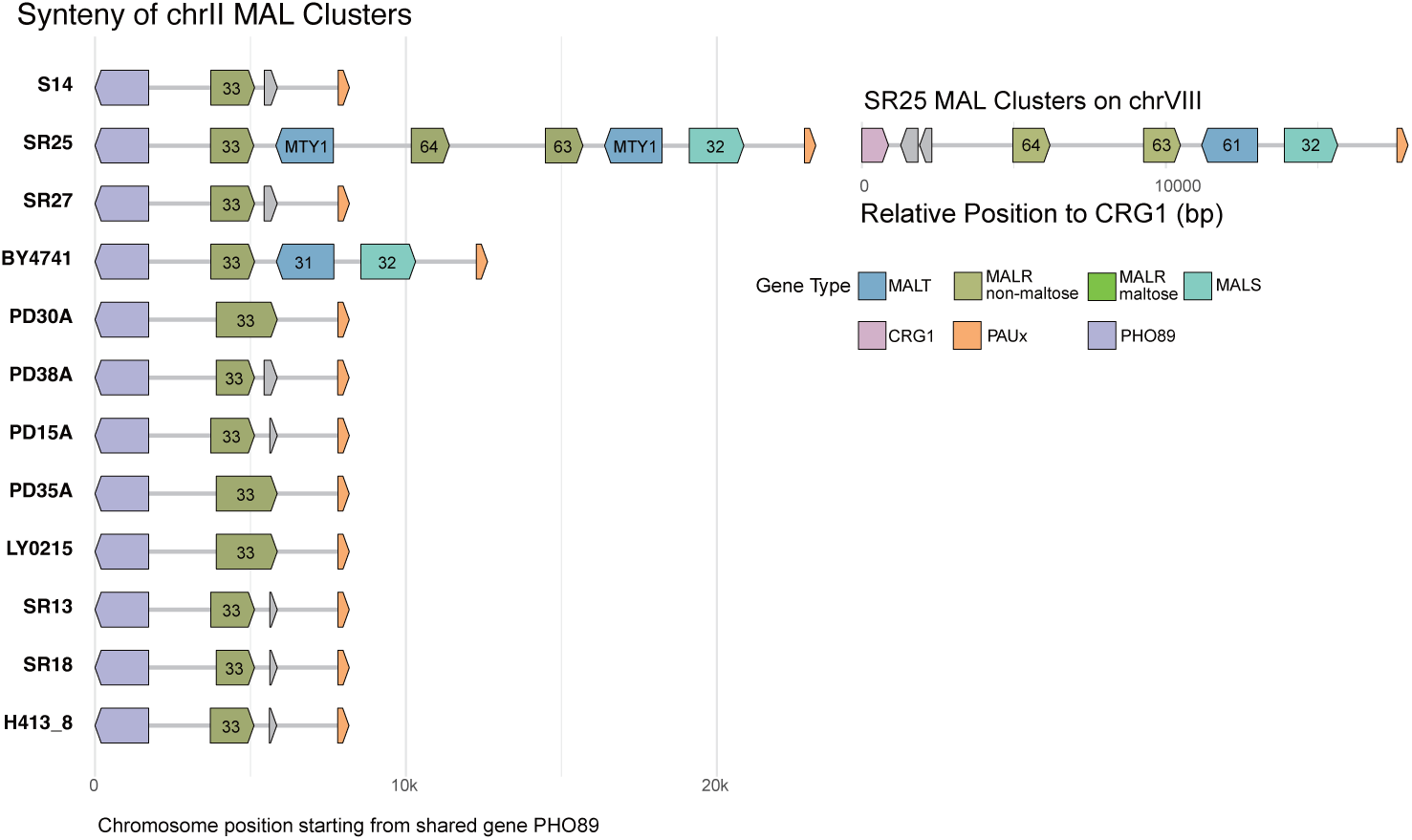
*MAL* loci synteny on chromosome II and VIII. Genes are colored by gene families and *MAL* genes are labeled by homology group.

**Supplementary Figure S6.**
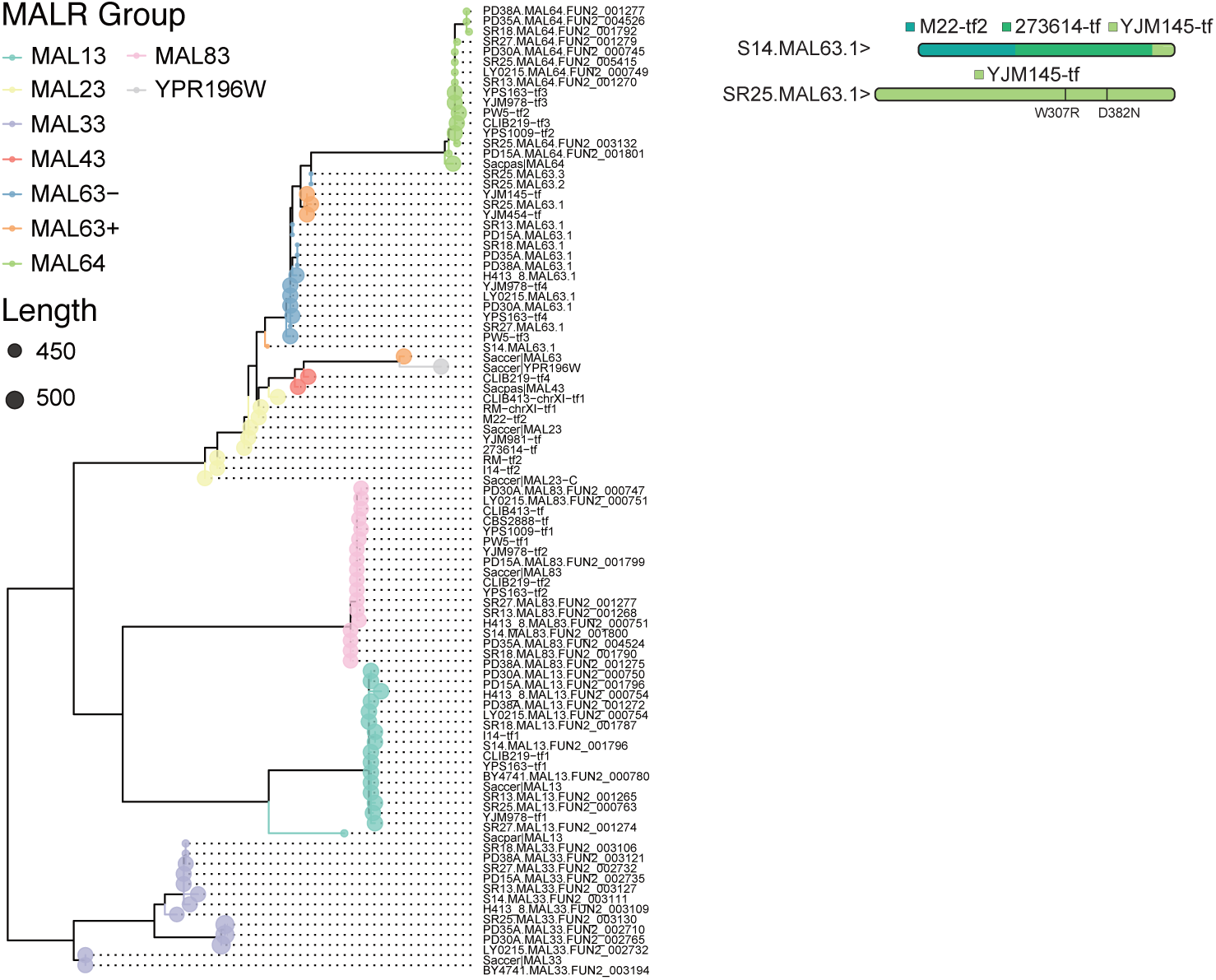
Grouping *MALR* by phylogeny and identity. **a.** *MALR* phylogenetic tree based on amino acid sequences. **b.** Shared alleles of *S14.MAL63.1* and *SR25.MAL63.1* with maltose-proficient *MALR*s (Weller et al. 2023).

**Supplementary Figure S7.**
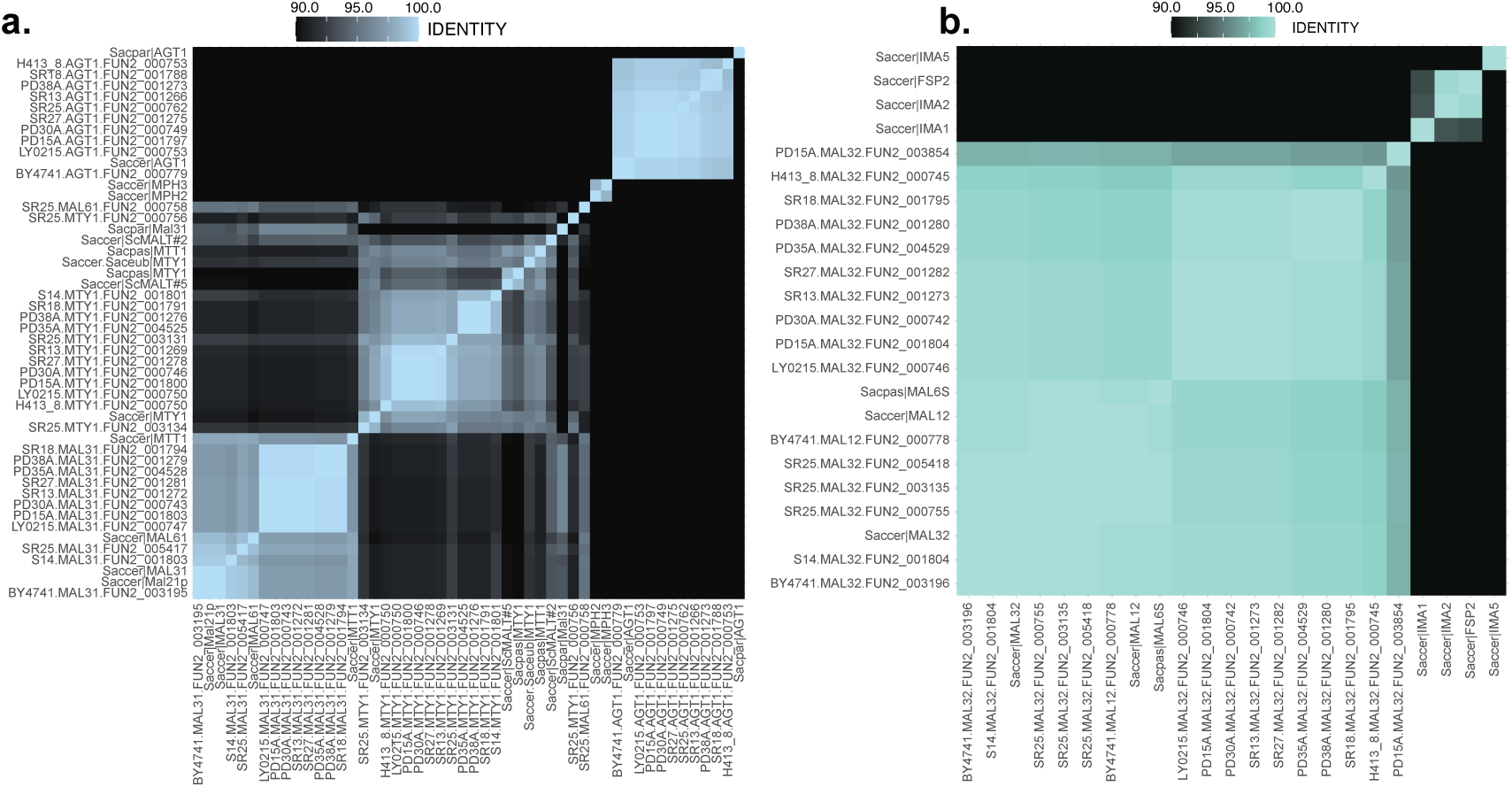
Amino acid sequence identity between **a.** *MALT* and **b.** *MALS*.

